# Preclinical characterization of mobocertinib highlights the putative therapeutic window of this novel EGFR inhibitor to EGFR exon 20 insertion mutations

**DOI:** 10.1101/2020.09.28.317560

**Authors:** Pedro E. N. S. Vasconcelos, Ikei S. Kobayashi, Susumu S. Kobayashi, Daniel B. Costa

## Abstract

**Background:** Epidermal growth factor receptor (EGFR) exon 20 insertion mutations account for 10% of all EGFR mutations and are mostly insensitive to approved EGFR-tyrosine kinase inhibitors (EGFR-TKIs). Novel EGFR-TKIs have been developed or repurposed for these mutants. A limited number of preclinical studies have detailed these EGFR-TKIs. We sought to use commercially available mobocertinib (TAK-788) to characterize the preclinical therapeutic window of this EGFR-TKI against EGFR mutations and to probe possible on-target mechanisms of resistance (EGFR-C797S).

**Methods:** We used models of EGFR mutations to probe representative 1^st^, 2^nd^, 3^rd^ generation, and in-development EGFR exon 20-active (poziotinib, mobocertinib) TKIs. We also introduced EGFR-C797S to these models to identify mechanisms of resistance.

**Results:** Cells driven by the most common EGFR exon 20 insertion mutations (A767_V769dupASV, D770_N771insSVD, H773_V774insH and others) were inhibited by in-development EGFR TKIs at doses below those affecting EGFR-wildtype; albeit more common EGFR mutations (exon 19 deletions and L858R) were inhibited more readily by mobocertinib and poziotinib. Mobocertinib was able to inhibit phosphorylation of EGFR in multiple preclinical models. The presence of EGFR-C797S led to >200-fold resistance in proliferation assays probing mobocertinib and osimertinib. Review of clinical studies of mobocertinib disclosed responses that could be lasting.

**Conclusions:** This is one of the initial reports to characterize the novel EGFR TKI mobocertinib and highlights its broad activity against EGFR mutants plus the therapeutic window to EGFR exon 20 insertion mutations; as well as EGFR-C797S as a possible mechanism of resistance. Further clinical development of mobocertinib merits continuation.

## Introduction

The majority of epidermal growth factor receptor (EGFR) and the structurally similar erb-b2 receptor tyrosine kinase 2 (ERBB2) exon 20 insertion mutations—representing 10% of all EGFR mutations—are unique among ErbB family members in that they activate the ATP binding pocket of these kinases without the typical conformational changes that enhance sensitivity to the majority of EGFR and/or ERBB2 tyrosine kinase inhibitors (TKIs) approved for clinical use in mid-2020 [1,2]. Over the last half a decade, novel EGFR- and/or ERBB2-TKIs or repurposed TKIs have been shown to have a slightly favorable therapeutic window for EGFR exon 20 insertion mutations in comparison to EGFR-wild type (WT) [3–6]. The ones that have reached clinical trial development for *EGFR* exon 20 insertion mutated non-small-cell lung cancer (NSCLC) include poziotinib, CLN-081, high dose osimertinib and mobocertinib; with differing levels of activity reported to date.

Mobocertinib, previously named AP32788 (ARIAD Pharmaceuticals) and subsequently TAK-788 (Takeda/Millennium Pharmaceuticals), was developed to have broad level of inhibition against EGFR kinase domain mutations based on the crystal structure of EGFR mutants and its covalent binding to EGFR [6,7]. The initial report of the preclinical activity of the drug was presented in abstract form in 2016 with basic assays disclosing lower inhibitory concentrations to a limited panel of EGFR and ERBB2 mutants, including exon 20 insertions, tested in comparison to EGFR-WT and ERBB2-WT proteins [6]. The initial clinical study of mobocertinib (ClinicalTrials.gov Identifier: NCT02716116) was initiated March 2016 and evolved to an extension cohort (named EXCLAIM) at the maximum tolerated dose of 160 mg a day [8]—the basis of the April 2020 Food and Drug Administration (FDA)’s breakthrough therapy designation of mobocertinib for the treatment of *EGFR* exon 20 insertion mutated NSCLC.

Our report sought to use commercially available mobocertinib to characterize the preclinical therapeutic window of this TKI against representative EGFR mutations and to probe possible on-target mechanisms of resistance (including EGFR-C797S); placing these results into context of the reported clinical activity of this drug and other in-development EGFR-TKIs.

## Methods

### Drugs

Erlotinib, afatinib, osimertinib (LC Laboratories), poziotinib (AdooQ BioScience) and mobocertinib (MedChemExpress) were dissolved in dimethyl sulfoxide and stored at −80°C.

### Preclinical models/cell lines

Ba/F3 (sequencing analysis was performed to confirm the presence of WT and mutant *EGFR*), BID007 (*EGFR*-A763_Y764insFQEA), BID019 (*EGFR*-N771_H772insH), PC-9 (*EGFR*-delE746_A750) and NCI-H1975 (H1975, *EGFR*-L858R+T790M) cell lines were maintained in RPMI 1640 medium (Mediatech) supplemented with 10% fetal bovine serum. In the case of EGFR-WT driven Ba/F3 cells, EGF 10 ng/mL was added to support growth. All cells were grown at 37°C in a humidified atmosphere with 5% CO_2_ and tested for lack of mycoplasma contamination (MycoAlert Mycoplasma Detection Kit, Lonza) prior to experiments (initiated within the initial 1-4 passages).

### Generation of compound EGFR-C797S mutations

The *EGFR*-C797S mutation was introduced into the *EGFR*-L858R sequence construct in the context of the MigR1 retrovirus vector (Addgene) using the QuickChange XL site-directed mutagenesis kit (Stratagene), as described by our group for other mutations [2].

### Proliferation assays

Cell viability was determined by CellTiter 96 aqueous one solution proliferation kit (Promega) and/or CellCountingKit-8 (CCK-8) (Dojindo Molecular Technologies) for Ba/F3 and other cells. Cells were plated in 96-well plates and then treated in the appropriate medium with or without EGFR TKIs for 3 days. Inhibitory proliferation curves and the 50% inhibitory concentration (IC_50_) were generated using GraphPad Prism (GraphPad Software).

### Protein-level analysis

Western blot lysates and preparation were performed as previously described [2,10]. Total EGFR, β-actin antibodies (Santa Cruz Biotechnology) and phospho-EGFR (pT1068) antibody (ThermoFisher) were diluted 1:1,000, while secondary antibodies were diluted 1:10,000.

## Results

### Preclinical evaluation of EGFR exon 20 insertion mutations against different classes of EGFR-TKIs in a Ba/F3 isogenic system

To highlight the potential therapeutic window of diverse classes of EGFR TKIs in different EGFR exon 20 insertion mutations, we selected 4 representative mutants (A763_Y764insFQEA, A767_V769dupASV, D770_N771insSVD and H773_V774insH) and more typical EGFR-TKI sensitive or resistant EGFR mutants (exon 19 deletion delL747_P753insS [del19], exon 21 L858R, del19+T790M and L858R+T790M) to contrast inhibitory concentrations with EGFR-WT (Fig.1A,B,C,D). The EGFR exon 20 insertion mutations that match the most frequent aberrations (A767_V769dupASV, D770_N771insSVD and H773_V774insH) in the Ba/F3 system displayed a favorable therapeutic window to poziotinib and mobocertinib (Fig.1A); while the therapeutic windows (IC_50_ EGFR-mutation divided by IC_50_ EGFR-WT) with erlotinib, afatinib and osimertinib were unfavorable (Fig.1A).

**Figure 1.**
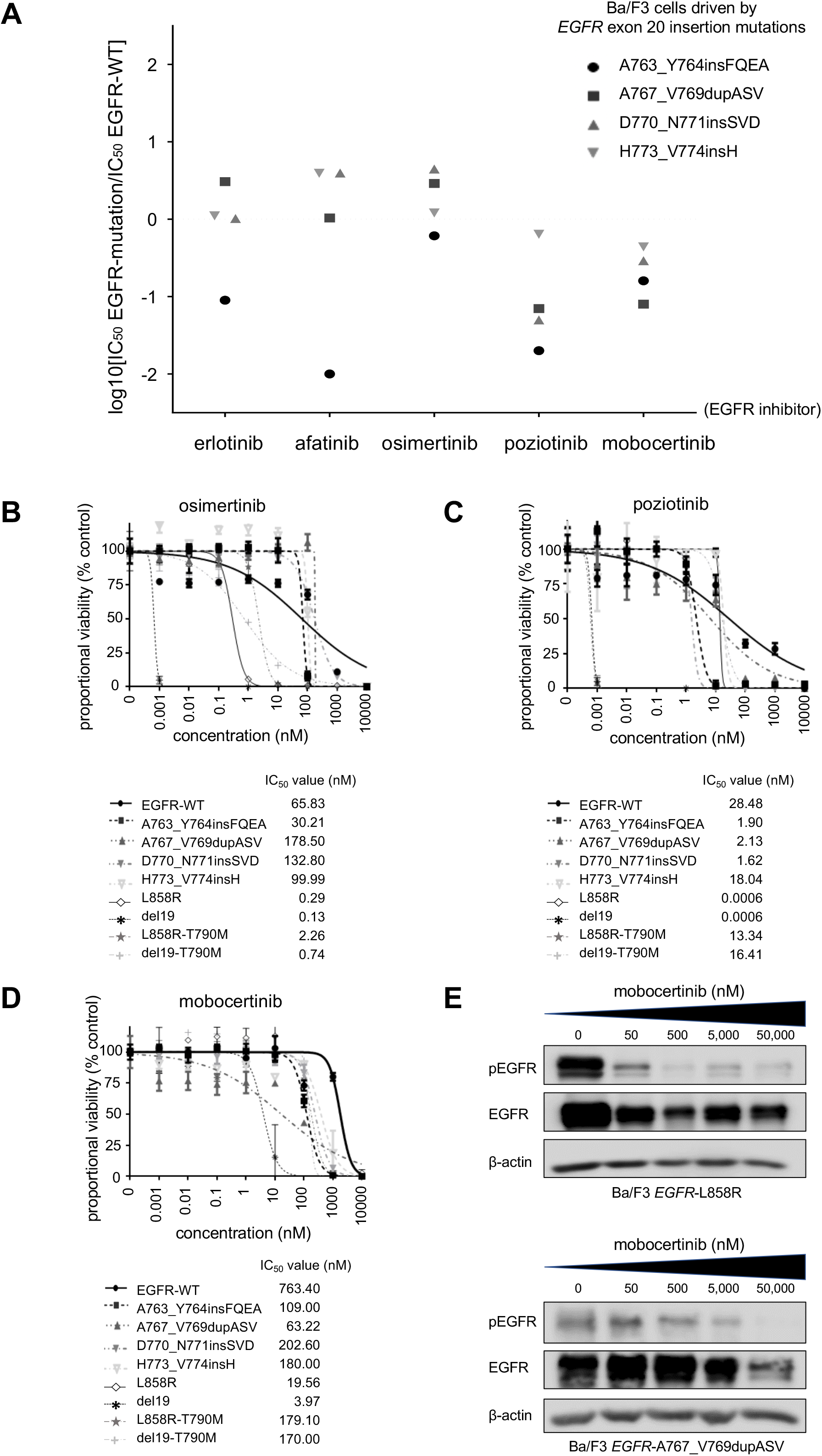
Ba/F3 system isogenic preclinical models of EGFR exon 20 insertions mutations to probe EGFR-TKIs, including mobocertinib. **A.** Therapeutic window of different EGFR-TKIs to a set of EGFR exon 20 mutations. Cells were plated at a density of 5000 cells per well (96-well plates) and grown over 3 days after treatment. Logarithm of the 50% inhibitory concentration (IC_50_) of EGFR exon 20 mutants compared to EGFR-WT is plotted (3 separate experiments are used to generate IC_50_). Values below 0 indicate sensitivity while values above 0 indicate resistance to EGFR-TKIs. **B.** Dose-response proliferation assays (proportional percent viability) of osimertinib for all tested EGFR mutants. **C.** Dose-response proliferation assays (proportional percent viability) of poziotinib for all tested EGFR mutants. **D.** Dose-response proliferation assays (proportional percent viability) of mobocertinib for all tested EGFR mutants. **E.** Western blotting of Ba/F3 cells driven by EGFR-L858R and EGFR-A767_V769dupASV. Cells were treated with the EGFR-TKI mobocertinib for 8 hours at the indicated ascending concentrations. pEGFR, phosphorylated EGFR at position 1068, total EGFR and β-actin as a loading control are displayed.

Erlotinib, afatinib (data not shown) were ineffective against compound EGFR-T790M bearing cells, poziotinib had values close to EGFR-WT (Fig.1C) while osimertinib (Fig.1B) and mobocertinib (Fig.1D) had favorable therapeutic windows. Of note, all the tested EGFR-TKIs had the lowest IC_50_ values for EGFR-del19 and -L858R bearing cells (Fig.1B,C,D). For mobocertinib, the values for IC_50_ were 3 to 51-fold higher for EGFR exon 20 insertions than del19 or L858R (Fig.1D) but lower than the IC_50_ of EGFR-WT (Fig.1A,D).

We also show that mobocertinib was able to induce, in a dose-dependent manner, inhibition of the phosphorylated state of EGFR in cells driven by EGFR-L858R and EGFR-A767_V769dupASV (Fig.1E); corroborating on-target engagement of this EGFR-TKI.

### Preclinical evaluation of mobocertinib against EGFR exon 20 insertion mutations in patient-derived lung cancer cell lines

To confirm the Ba/F3 preclinical model findings, we used a second set of preclinical systems with patient-derived lung adenocarcinoma cell lines. The *EGFR*-A763_Y764insFQEA-bearing BID007 cells and the *EGFR*-N771_H772insH-bearing BID019 cells were inhibited by mobocertinib in a dose-dependent manner (Fig.2A) similar to the Ba/F3 counterpart results (Fig.1). Cell lines harboring the EGFR-TKI sensitive *EGFR*-delE746_A750 mutation (PC-9) were the most sensitive while lines harboring the erlotinib/afatinib/poziotinib resistant *EGFR*-L858R+T790M mutation (H1975) were the least sensitive (Fig.2A).

**Figure 2.**
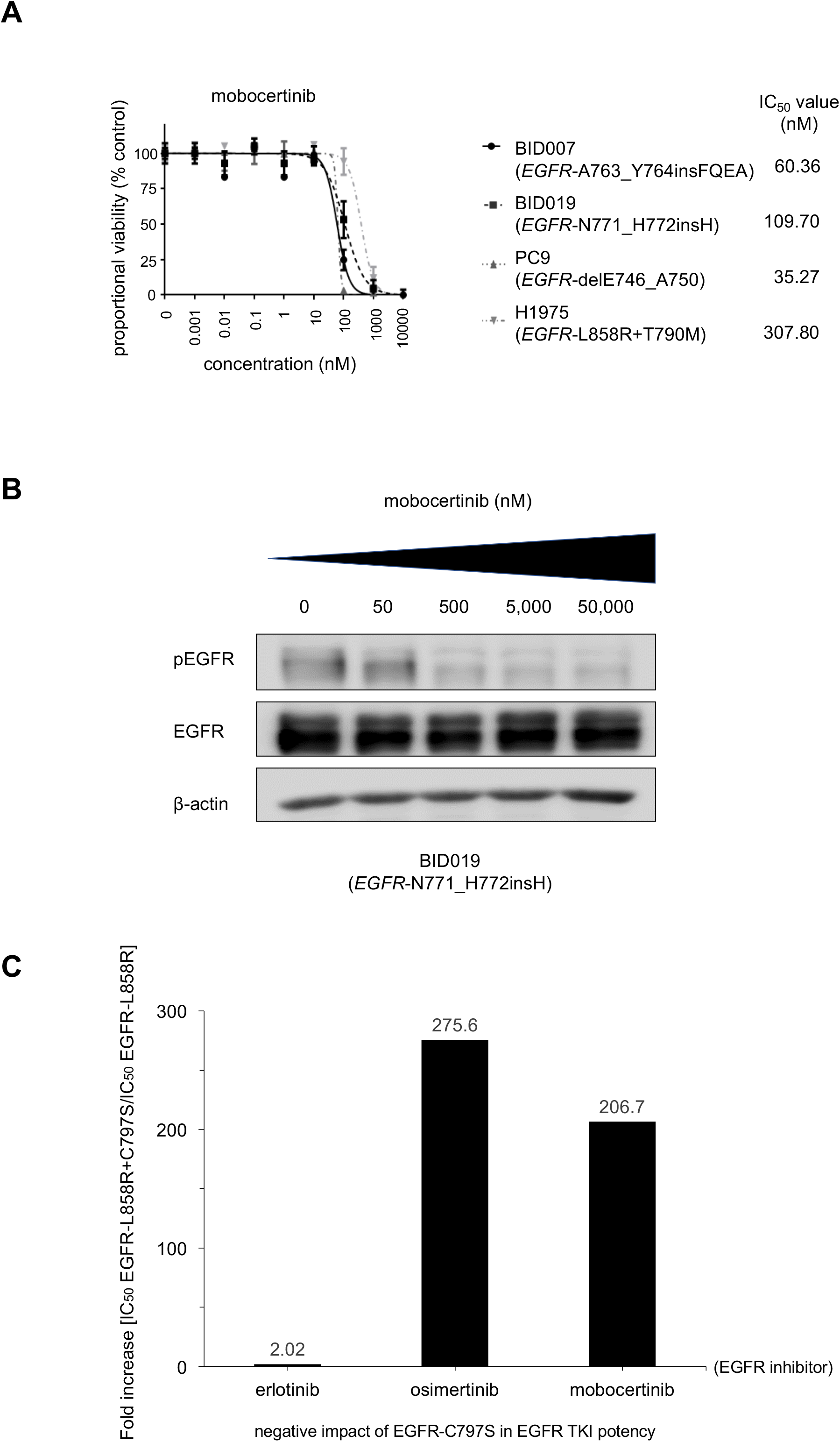
Patient-derived cancer cell line preclinical models of EGFR exon 20 insertions mutations to probe mobocertinib and the role of *EGFR*-C797S as a mechanism of resistance. **A.** Dose-response proliferation assays (proportional percent viability) of mobocertinib for all tested patient-derived *EGFR* mutated lung adenocarcinoma cell lines. The main driver *EGFR* mutation is indicated for each cell. **B.** Western blotting of cell line BID019 (*EGFR*-N771_H772insH). Cells were treated with the EGFR-TKI mobocertinib for 8 hours at the indicated ascending concentrations. pEGFR, phosphorylated EGFR at position 1068, total EGFR and β-actin as a loading control are displayed. **C.** Relative fold increase in the 50% inhibitory concentration (IC_50_) of EGFR-L858R+C797S compared to EGFR-L858R in the Ba/F3 system. 3 separate experiments are used to generate IC_50_ for each EGFR-TKI tested, erlotinib, osimertinib and mobocertinib.

We also subjected BID019 to a dose-response exposure experiment to evaluate intracellular signaling, and observed dose-dependent inhibition of phosphorylated EGFR (Fig. 2B) in this model.

### Role of EGFR-C797S as an on-target mechanism of resistance to mobocertinib

Mobocertinib is an irreversible EGFR inhibitor designed to covalently bind to amino acid cysteine at position 797 of the EGFR’s kinase domain [6,7]. Therefore, we generated a system harboring a mobocertinib-sensitive *EGFR* mutation *in cis* to the *EGFR*-C797S mutation (Fig.1C). The compound EGFR-C797S mutated protein product negated the inhibitory ability of mobocertinib and osimertinib, as indicated by the >200-fold shift in measured IC_50_ values (Fig.1C). Erlotinib, a reversible EGFR-TKI that does not bind covalently to EGFR-C797, was minimally disturbed by the presence of the compound EGFR-C797S product (Fig.1C).

## Discussion

The clinical care of advanced NSCLC in 2020 is anchored on tumor-based genomic and immune biomarkers. EGFR was the first “targetable” driver oncogene and perhaps one of the most clinically significant; however, EGFR-TKIs are only approved for tumors harboring *EGFR*-del19, -L858R, -L861Q, G719X, -S768I and -T790M. Off label use of approved EGFR TKIs is further supported for EGFR-exon 18 indels/E709X, -exon 19 insertions, -exon 20 A763_Y764insFQEA, exon 18-25 kinase domain duplications and - rearrangements [1,2,5,9,10]. The identification of EGFR-TKIs or other therapies with a favorable therapeutic window for patients with *EGFR* exon 20 insertion mutated tumors continues to be a significant unmet necessity.

We characterize the irreversible EGFR-TKI mobocertinib in the current study. Mobocertinib was generated using a structure-based design to increase potency and selectivity to EGFR mutations in relation to EGFR-WT [6,7]. It has a kinase selectivity profile for EGFR, ERBB2 and other ErbB family members with minimal selectivity against other common kinases [7]. Initial studies disclosed broad level of activity against most types of common EGFR mutations, including exon 20 insertion mutations, in isogeneic cell lines and *in vivo* murine models [6]. Our group was able to show that indeed mobocertinib is a pan-active EGFR-TKI with a favorable therapeutic window (inhibitory profiles below those of EGFR-WT) for all EGFR mutations tested (Fig.1,Fig.2). It is most potent against the canonical EGFR mutations, del19 and L858R (Fig.1D,Fig.2A), but still selective—albeit with a narrower therapeutic window—against the most common EGFR exon 20 insertion mutations, A767_V769dupASV, D770_N771insSVD and H773_V774insH (Fig.1A). We confirmed this selectivity potential using isogenic cell line models (Fig.1), NSCLC cancer cell lines with *EGFR* exon 20 insertions (Fig.2) and at the intracellular signaling level (Fig.1E,Fig.2B). Our data, to the best of our knowledge, shows for the first time that mobocertinib is susceptible to on-target resistance mediated by the *EGFR*-C797S mutation that negates the drug’s covalent binding cysteine; in a pattern also seen with the 3^rd^ generation first-in-class EGFR-TKI osimertinib (Fig.2C). These results form a sound preclinical complementary basis to support the clinical results of mobocertinib. The most updated results of the initial phase I/II and extension cohort (EXCLAIM) trial of mobocertinib (NCT02716116) were presented in September 2020 (Table1). The recommended phase 2 dose was established at 160 mg/day and 28 patients were treated in the expansion cohort at this initial dose [8]. The overall response rate (ORR) was 43% (95%CI:24%-63%), with a disease control rate of 86% (95%CI:67%-96%) and median progression free survival (PFS) of 7.3 months (95%CI:4.4-15.6). Toxicities were noted in the majority of cases and were mostly related to gastrointestinal events (i.e., diarrhea, nausea) and skin rash [8] that are markers of EGFR-WT adverse events common to most EGFR-TKIs. The narrow therapeutic window observed in our preclinical report (Fig.1D) substantiate the pattern of adverse events seen in the clinical study. The drug’s observed activity led to a FDA breakthrough therapy designation of mobocertinib for the treatment of *EGFR* exon 20 insertion mutated NSCLC in April 2020. These initial results also spawned a randomized phase III trial of mobocertinib versus standard platinum-doublet chemotherapy in 318 treatment-naïve advanced NSCLC with *EGFR* exon 20 insertion mutations (EXCLAIM-2, NCT04129502) that commenced in 2019. The results of the latter trial may help establish the evidence-based 1^st^ line treatment strategy for *EGFR* exon 20 mutated NSCLC in the decade to come.

**Table 1.** Summary of the outcomes from clinical trials of select EGFR tyrosine kinase inhibitors (EGFR-TKIs) for *EGFR* exon 20 insertion mutated lung cancer, including mobocertinib, poziotinib and high dose osimertinib. Footnote: ORR, overall response rate; DCR, disease control rate; PFS, progression-free survival (median); CI, confidence interval. Data was based on references [9] (mobocertinib), [12] (poziotinib) and [13] (osimertinib).

| EGFR TKI/<br>starting dose/<br>number cases | trial name/<br>NCT no./<br>reference | ORR | DCR | PFS |
| --- | --- | --- | --- | --- |
| osimertinib<br>160 mg/day<br>(n=21) | ECOG-ACRIN 5162<br>NCT03191149<br>ASCO 2020 [13] | 24% | 82% | 9.6 months<br>(95%CI 4.1,10.7 m.) |
| poziotinib<br>16 mg/day<br>(n=115) | ZENITH20<br>NCT03318939<br>ASCO 2020 [12] | 14.8%<br>(95%CI 8.9,22.6%) | 68.7%<br>(95%CI 59.4,77.0%) | 4.2 months<br>(95%CI 3.7,6.8 m.) |
| mobocertinib<br>160 mg/day<br>(n=28) | EXCLAIM<br>NCT02716116<br>ESMO 2020 [9] | 43%<br>(95%CI 24,63%) | 86%<br>(95%CI 67,96%) | 7.3 months<br>(95%CI 4.4,15.6 m.) |

Other EGFR-TKIs currently in clinical development—such as poziotinib and CLN-081—have also shown a reproducible therapeutic window in preclinical models [4,5,10]. The clinical trial of poziotinib for *EGFR* exon 20 insertion mutated NSCLC (ZENITH20, NCT03318939) has led to suboptimal initial efficacy results, with ORR below 15% and median PFS below 5 months (Table1)—results that may be explained by dose limiting skin and gastrointestinal adverse events at the 16 mg/day dosing [11]. Further development of alternative dosing schemes of poziotinib are ongoing (NCT03318939). The initial dose escalation of CLN-081 [5] is ongoing with some initial responses noted in the target cohort (NCT04036682). Another EGFR-TKI strategy based on our group’s preclinical models [3] is the use of a higher than standard dose of osimertinib. The ECOG-ACRIN 5162 trial (NCT03191149) showed the dosing scheme of osimertinib 160 mg/day (double the FDA approved dose) was tolerable but only induced responses in the minority of cases [12], with a ORR below 25% (Table1). Perhaps higher doses of osimertinib may be necessary in future studies. Of note, poziotinib and mobocertinib are also undergoing clinical trial evaluation for *ERBB2* exon 20 insertion mutated NSCLC in the aforementioned clinical studies. The ERBB2-TKI pyrotinib (NCT02535507) was recently shown to have a modest ORR of 30% (95%CI:18.8%-43.2%) and median PFS of 6.9 months (95%CI:5.5-8.3) in *ERBB2* mutated NSCLC [13]. It is likely other EGFR and/or ERBB2-TKIs will enter this burgeoning clinical trial space. Other strategies to target these tumors include antibody-drug conjugates (ADCs). The most robust results have been shown for ERBB2 ADCs, with ado-trastuzumab emtansine and trastuzumab deruxtecan showing efficacy and safety in NSCLC [14]. The dual EGFR-MET ADC amivantamab has been tested in *EGFR* mutated NSCLCs (CHRYSALIS, NCT02609776), and initial results in 39 patients with *EGFR* exon 20 insertion mutated NSCLC [15] showed ORR of 36% (95%CI:21%-53%) and median PFS of 8.3 months (95%CI:3.0-14.8)—the basis of a FDA breakthrough therapy designation in May 2020.

In summary, our preclinical results and the evolving clinical trial program of mobocertinib assert this pan-active EGFR-TKI as a promising treatment strategy for *EGFR* exon 20 insertion mutated NSCLC. However, the same data highlight the potential for high rates of EGFR-WT mediated adverse events and susceptibility to *EGFR*-C797S as a mechanism of resistance. The robust pipeline of EGFR-TKIs and ADCs with preclinical and early clinical activity against *EGFR* and *ERBB2* exon 20 insertion mutated tumors hold the promise of a much-needed drug approval for these important cohorts of NSCLC.

## Acknowledgements/Funding

This work was funded in part through National Institutes of Health (NIH)/National Cancer Institute (NCI) grants R37 CA218707 (to D. B. Costa), R01 CA240257 (to S. S. Kobayashi) plus Department of Defense LC170223 (to S. S. Kobayashi)

## Notes

*Disclosure of Potential Conflicts of interest:* DBC reports personal fees (consulting fees and honoraria) and nonfinancial support (institutional research support) from Takeda/Millennium Pharmaceuticals, AstraZeneca, Pfizer and BluePrint Medicine, as well as nonfinancial support (institutional research support) from Merck Sharp and Dohme Corporation, Merrimack Pharmaceuticals, Bristol-Myers Squibb, Clovis Oncology, Spectrum Pharmaceuticals, Tesaro, all outside the submitted work. SSK reports research support from Boehringer Ingelheim, MiNA Therapeutics, and Taiho Therapeutics, as well as personal fees (honoraria) from Boehringer Ingelheim, Bristol Meyers Squibb, and Takeda Pharmaceuticals; outside the submitted work. No other conflict of interest is reported.

### Competing Interest Statement

DBC reports personal fees (consulting fees and honoraria) and nonfinancial support (institutional research support) from Takeda/Millennium Pharmaceuticals, AstraZeneca, Pfizer and BluePrint Medicine, as well as nonfinancial support (institutional research support) from Merck Sharp and Dohme Corporation, Merrimack Pharmaceuticals, Bristol-Myers Squibb, Clovis Oncology, Spectrum Pharmaceuticals, Tesaro, all outside the submitted work. SSK reports research support from Boehringer Ingelheim, MiNA Therapeutics, and Taiho Therapeutics, as well as personal fees (honoraria) from Boehringer Ingelheim, Bristol Meyers Squibb, and Takeda Pharmaceuticals; outside the submitted work. No other conflict of interest is reported.

